# Site-directed crosslinking identifies the stator-rotor interaction surfaces in a hybrid bacterial flagellar motor

**DOI:** 10.1101/2021.01.07.425829

**Authors:** Hiroyuki Terashima, Seiji Kojima, Michio Homma

## Abstract

The bacterial flagellum is the motility organelle powered by a rotary motor. The rotor and stator elements of the motor are embedded in the cytoplasmic membrane. The stator units assemble around the rotor, and an ion flux (typically H^+^ or Na^+^) conducted through a channel of the stator induces conformational changes that generate rotor torque. Electrostatic interactions between the stator protein PomA in *Vibrio* (MotA in *Escherichia coli*) and the rotor protein FliG have been suggested by genetic analyses, but have not been demonstrated directly. Here, we used site-directed photo- and disulfide-crosslinking to provide direct evidence for the interaction. We introduced a UV-reactive amino acid, *p*-benzoyl-L-phenylalanine (*p*BPA), into the cytoplasmic region of PomA or the C-terminal region of FliG in intact cells. After UV irradiation, *p*BPA inserted at a number of positions formed a crosslink with FliG. PomA residue K89 gave the highest yield of crosslinks, suggesting that it is the PomA residue nearest to FliG. UV-induced crosslinking stopped motor rotation, and the isolated hook-basal body contained the crosslinked products. *p*BPA inserted to replace residues R281 or D288 in FliG formed crosslinks with the *Escherichia coli* stator protein, MotA. A cysteine residue introduced in place of PomA K89 formed disulfide crosslinks with cysteine inserted in place of FliG residues R281 and D288, and some other flanking positions. These results provide the first demonstration of direct physical interaction between specific residues in FliG and PomA/MotA.

## Importance

The bacterial flagellum is a unique organelle that functions as a rotary motor. The interaction between the stator and rotor is indispensable for stator assembly into the motor and the generation of motor torque. However, the interface of the stator-rotor interaction has only been defined indirectly by mutational analysis. Here, we detected the stator-rotor interaction using site-directed photo- and disulfide-crosslinking approaches. We identified several residues in the PomA stator, especially K89, that are in close proximity to the rotor. Moreover, we identified several pairs of stator and rotor residues that interact. This study directly demonstrates the nature of the stator-rotor interaction and suggests how stator units assemble around the rotor and generate torque in the bacterial flagellar motor.

## Introduction

F-type ATP synthase, V/A-type ATPase, and the bacterial flagellum are well-known examples of ion-driven molecular rotary motors (1, 2). The flagellum of bacteria other than spirochetes has a helical filament that extends from the cell surface and functions as a rotary screw to propel swimming. The rotary motor of the bacterial flagellum consists of a rotor surrounded by varying numbers of stator units, with both the rotor and stator embedded in the cytoplasmic membrane (3-6). The rotor contains a transmembrane MS-ring and an attached cytoplasmic C-ring below the MS-ring (7, 8). In many bacteria, the C-ring contains the three proteins FliG, FliM and FliN. Mutations in the genes encoding these proteins can confer *fla, mot* and *che* phenotypes, corresponding to deficiencies in flagellar formation, motor rotation, or the switching between CCW and CW rotation (5, 9). FliG is thought to interact with the stator units to generate torque (10, 11).

MotA and MotB in *Escherichia coli* (*E. coli*), and PomA and PomB in *Vibrio* species are the membrane proteins that comprise the stator complex (12-16). The A subunit has four transmembrane segments (TM) and a large cytoplasmic region between TM2 and TM3. The B subunit has one TM in its N-terminal region and a peptidoglycan-binding (PGB) domain in its C-terminal region (17-19). In *E. coli*, at least 11 stator units can assemble around, and interact with, the rotor. They are anchored at the proper position by the PGB domain, and once incorporated, the stator unit is activated for ion conduction and motor rotation (20-25). The coupling ion is a proton in the *E. coli* motor and a sodium ion in the *Vibrio* motor; it is conducted to the cytoplasm through an ion-transporting pathway in the stator complex (26, 27). A conserved aspartate residue in the TM of the B subunit receives the coupling ion from outside the cytoplasm, and the ion then dissociates into the cytoplasm (28-30). The ion-binding and release cycle induces conformational changes in the stator complex that change the interactions between the A subunit of the stator and FliG of the rotor (31, 32).

Earlier biochemical studies demonstrated the stator-rotor interaction using a His-tag pull-down assay (33). Stator interactions with the rotor protein have been examined in detail in *E. coli* using genetic analysis (34-36). MotA and FliG in *E. coli* have well-conserved charged residues, R90 and E98 in MotA and R281, D288 and D289 in the C-terminal domain of FliG (Fig. S1A, S1B). Charge neutralization or inversion of these residues leads to defects in motility, and the proper combinations of charge reversals between MotA and FliG synergistically rescue motility. The conserved charged residues in the A subunit are important for torque generation and assembly of the stator units into the motor (24, 25, 37). In contrast, charge neutralization or inversion of the corresponding residues in *Vibrio* did not abolish motility, suggesting that the charged residues are important, but not critical, for flagellar rotation in *Vibrio* (Fig. S1A, S1B) (38, 39). This result implies that additional residues contribute to motor rotation. Since the stator-rotor interaction in the *Vibrio* flagellar motor is likely to be more extensive than in *E. coli* (25, 40), *Vibrio* PomA is better suited for the examination of interactions between the stator A subunit and FliG of the rotor.

Structural information is indispensable for understanding the mechanism that produces rotation of the bacterial flagellar motor. Until recently, we had only low-resolution density maps of the stator unit obtained through single-particle analysis using electron microscopy (41, 42). However, atomic resolution structures of MotA/MotB from *Campylobacter jejuni, Clostridium sporogenes, Bacillus subtilis* and other species have been reported recently (43, 44). The structures resemble the structure of ExbB (45-47).

MotA and PomA share a weak sequence homology with ExbB of the Ton bacterial transport system, which transports relatively large molecules, such as siderophores and vitamin B12, into the cell (43, 44). The MotA/MotB and PomA/PomB complexes exist as a 5:2 hetero-heptamer, although the stoichiometry was previously proposed as a 4:2 hetero-hexamer (14, 16, 42). The atomic resolution structures of the stator provide insight into its organization and its contribution to flagellar rotation. Dynamic interactions between the stator and rotor generate torque that rotates the flagellum. However, the molecular details of the stator-rotor interaction remain obscure.

In this study, we probed residues of PomA for the ability to crosslink with FliG using a site-directed *in vivo* photo-crosslinking technique. This technique allows *p*BPA, a phenylalanine derivative containing a UV-reactive benzophenone group, to be charged to an amber suppressor tRNA *in vivo* by a mutated tyrosyl-tRNA synthase from *Methanococcus jannaschii. p*BPA can be incorporated into any protein of interest by introducing an amber codon into the target position (48). The *p*BPA incorporated into the protein forms a covalent bond with a close C-H bond upon UV irradiation. Another approach utilizes disulfide bond formation between cysteine residues inserted at desired positions in two interacting proteins. Here, we report that photo-crosslinked and disulfide-crosslinked products are formed between targeted residues of PomA and FliG. This work provides the first direct evidence to show which residues in the stator and rotor are in close juxtaposition and suggests interactions that are responsible for stator assembly and torque generation in the flagellar motor.

## Results

### Effect of PomA with pBPA on E. coli cell motility

Because we were unable to adapt the technique for introducing *p*BPA into *Vibrio*, we performed a photo-crosslinking experiment in *E. coli* using a chimeric PomA/PotB stator unit that functions in *E. coli*. PotB is a chimeric protein in which the N-terminal region of PomB is fused to the C-terminal region of *E. coli* MotB (49). We introduced *p*BPA at each position in PomA from E74 to F104. This region contains the important charged residues, R88 and E96, which are proposed to interact with FliG (Fig. S1A, S1C). First, we examined whether PomA with the *p*BPA insertions confers motility to an *E. coli* Δ*motAB* null strain. The *p*BPA substitutions at L76, I77, I80, A84, G91, L95, N102 and F104 caused loss of motility (Fig. S2). None of these proteins other than the one with a substitution at L95 could be detected in the cells (Fig. S3). PomA could be detected in all the motile cells (Fig. S3). PomA with substitutions at A87, R88 and E96 support much less motility than wild-type PomA (Fig. S2). These results suggest that residues A87, R88, L95 and E96 are at, or are very close to, the sites that are important for motor function.

### Detection of photo-crosslinked products of PomA and FliG

We UV-irradiated cells expressing PomA containing *p*BPA and then probed by immunoblotting with anti-FliG antibody for crosslinked products. Crosslinking was observed when *p*BPA replaced the residues D85, R88, K89, G90, F92, L93 and E96, whereas it was not observed in the vector control or with wild-type PomA (Fig. 1A lower panels; for the complete set of mutants, see Fig. S3). This result indicates that PomA with *p*BPA formed photo-crosslinks with FliG. The only crosslinked product identified by probing with anti-PomA antibody was observed when *p*BPA replaced K89, presumably because of the low titer of the anti-PomA antibody (Fig. 1A, upper panels). Consistent with this result, the signal intensity of crosslinked products detected by the anti-FliG antibody was strongest when *p*BPA replaced K89 (Fig. 1A).

**Fig. 1.**
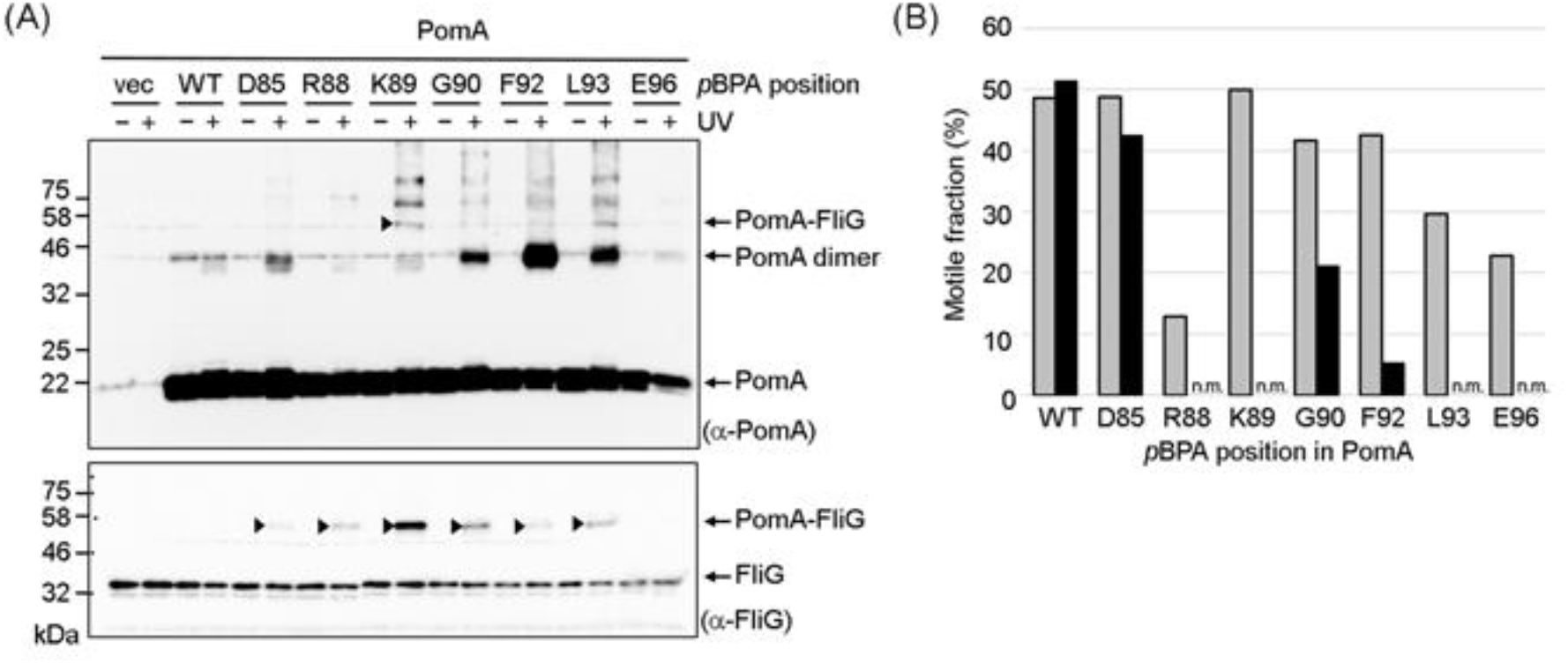
Photo-crosslinking between plasmid-encoded *Vibrio* PomA and endogenous *E. coli* FliG. (A) *Vibrio* PomA and chimeric PotB were expressed from plasmid pYS3, and the amber suppressor tRNA and the mutated tyrosyl-tRNA synthase were expressed from plasmid pEVOL-pBpF in the *E. coli* Δ*motAB* strain, RP6894. After photo-crosslinking, whole-cell lysates were prepared and analyzed using immunoblotting. The upper and lower panels show immunoblot images produced with anti-PomA and anti-FliG antibodies, respectively. The crosslinked products are indicated by black arrowheads. We showed the image of the photo-crosslinked product of PomA E96*p*BPA in Fig. S3C because we could not detect it in this immunoblot. Bands with higher molecular weight were derived from non-specific crosslinking between stator units or non-specific crosslinking of PomA with other proteins. (B) The motile fraction of *E. coli* RP6894 cells expressing PomA/PotB before and after UV irradiation in free-swimming. The gray box shows the motile fraction before UV irradiation. The black box shows the motile fraction after UV irradiation. At least 30 freely suspended cells were analyzed for each mutant with dark-field microscopy. Abbreviations: n.m., nonmotile.

We determined the fraction of motile cells before and after UV irradiation because we expected that the PomA-FliG crosslink would block motor function. With wild-type PomA, almost the same fraction of cells swam before and after UV irradiation.

Irradiation of the cells containing *p*BPA replacing R88, K89, L93 and E96 did not swim after UV irradiation, and cells with *p*BPA replacing G90 and F92 mutants had a lower motile fraction (Fig. 1B). These results suggest that the residues R88, K89, G90, F92, L93 and E96 of PomA are close enough to the rotor C-ring to form crosslinks with FliG.

### pBPA-labeled PomA binds to the C-ring after photo-crosslinking

Most of the photo-crosslinked products described above could have arisen through crosslinking between PomA freely diffusing in the cytoplasmic membrane and FliG freely diffusing in the cytoplasm (please see Fig. S4, as described below). Therefore, we examined whether photo-crosslinked PomA was associated with the isolated hook-basal body (HBB) fraction. After UV irradiation of cells expressing PomA with *p*BPA replacing D85, R88, K89, G90, F92, L93 or E96, the cells were solubilized using TritonX-100, and then HBBs were isolated using ultra-centrifugation. In immunoblots developed using the anti-FliG antibody, the HBB fractions contained photo-crosslinked products when *p*BPA replaced D85, R88, K89, and L93 (Fig. 2). The crosslinked products were most evident when *p*BPA replaced PomA K89. Unfortunately, we could not see PomA/PotB bound to the C-ring in electron micrographs. These results confirm that the PomA/PotB complex associates with FliG assembled into the rotor and support the suggestion that that K89 is in the closest proximity to FliG.

**Fig. 2.**
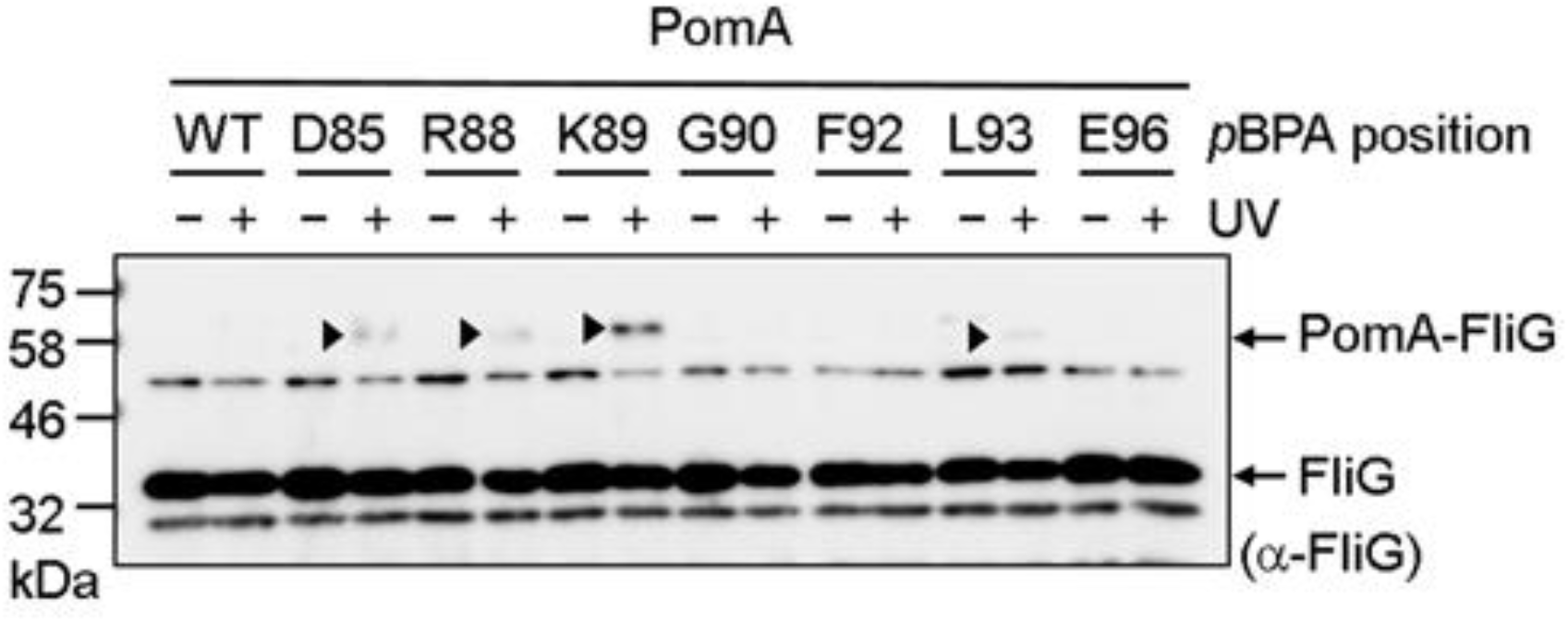
Photo-crosslinking between plasmid-borne *Vibrio* PomA and endogenous *E. coli* FliG in the isolated hook-basal body. *Vibrio* PomA and chimeric PotB were expressed from plasmid pYS3, and the amber suppressor tRNA and the mutated tyrosyl-tRNA synthase were expressed from plasmid pEVOL-pBpF in the *E. coli* Δ*motAB* strain RP6894. The panel shows immunoblot images visualized with an anti-FliG antibody. The crosslinked products are indicated by black arrowheads.

### pBPA-labeled PomA binds to FliG diffusing freely in the cytoplasm

We next examined whether PomA/PotB interacts with freely diffusing FliG. We expressed PomA with *p*BPA replacing D85, R88, K89, G90, F92, L93 or E96 and PotB from plasmid pTSK170 in *E. coli* Δ*flhDC* cells, which lack all flagellar proteins. We co-expressed *E. coli* FliG in these cells. After UV irradiation of these cells, we probed the photo-crosslinked products by immunoblotting with anti-PomA and anti-FliG antibodies. PomA with *p*BPA replacing K89 produced a large amount of crosslinked product, whereas the other proteins showed fewer crosslinked products (Fig. S4). We speculate that photo-crosslinked products could be detected in this experiment using the anti-PomA antibody because we produced FliG in great excess.

### FliG residues that interact with PomA

Residues R281 and D288 of *E. coli* FliG have been implicated in interacting with MotA. First, we investigated whether FliG with *p*BPA introduced at these two positions formed crosslinked products with endogenous *E. coli* MotA (Fig. 3, S1B). Indeed, when these proteins were expressed as the sole FliG, we detected crosslinked FliG-MotA using anti-MotA antibody, whereas the vector control and wild-type FliG did not form the crosslinked product. Further, FliG with *p*BPA replacing K264, D289 and R297 also did not form the crosslinked products. This result suggests that R281 and D288 are in close proximity to MotA.

**Fig. 3.**
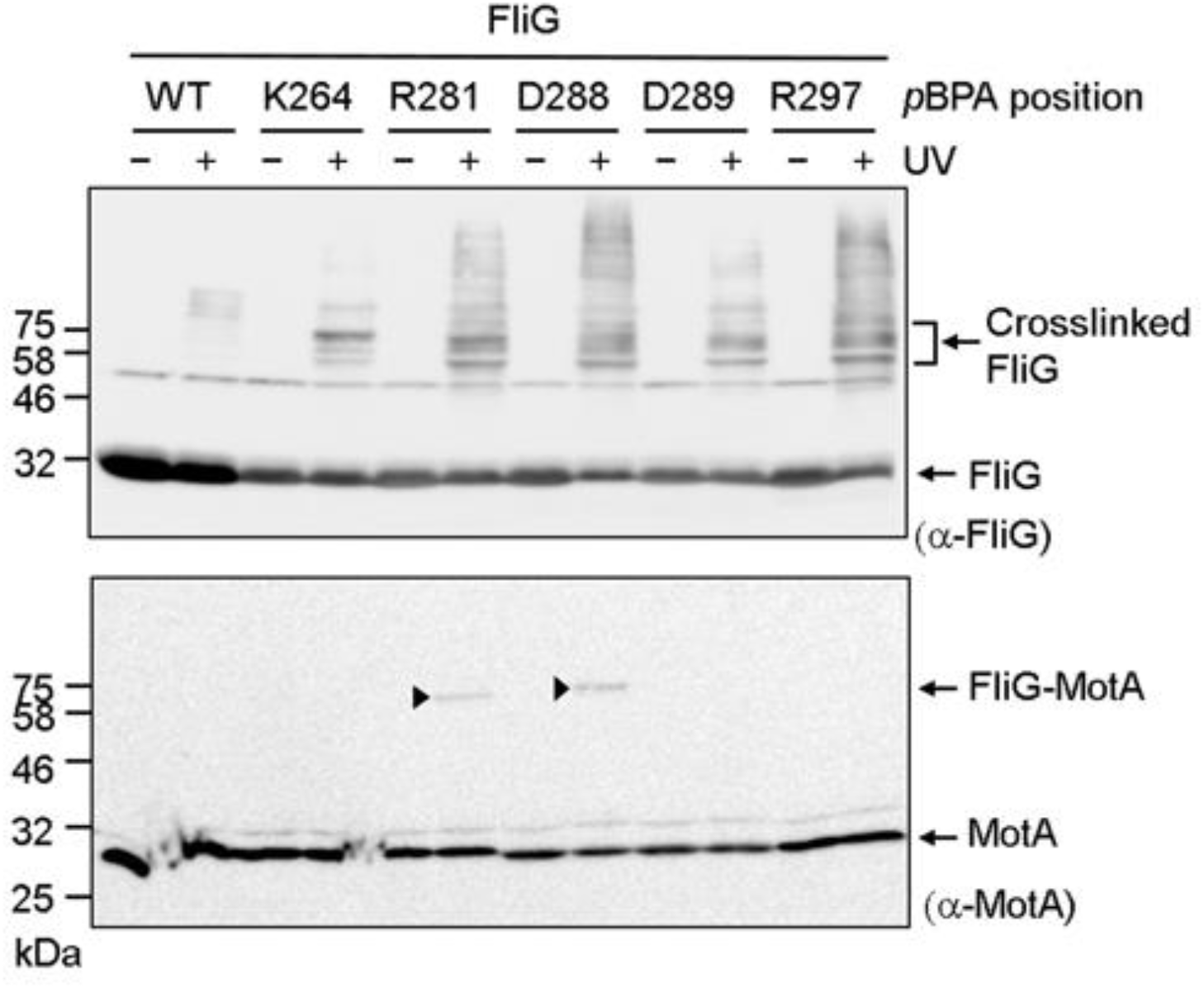
Photo-crosslinking between plasmid-borne *E. coli* FliG and endogenous *E. coli* MotA. *E. coli* FliG was expressed from plasmid pTY801, and the amber suppressor tRNA and the mutated tyrosyl-tRNA synthase were expressed from plasmid pEVOL-pBpF in the *E. coli* Δ*fliG* strain DFB225. The upper and lower panels show immunoblot images made using anti-MotA and anti-*E. coli* FliG antibodies, respectively. The crosslinked products are indicated by black arrowheads. Bands with higher molecular weight in the immunoblot with anti-FliG antibody were derived from non-specific interactions of FliG with other proteins.

Next, we investigated whether PomA containing cysteine replacements for residues K89 and L93 could form disulfide crosslinks with FliG containing cysteine replacements at critical residues. Previous genetic studies (25) suggested that PomA K89 interacts with residues R301, D308 and D309 of *Vibrio* FliG. Therefore, we co-expressed PomA K89C/PotB together with the Q280C, R281C, A282C, D288C or D289C variants of *E. coli* FliG in a Δ*motA/*Δ*fliG E. coli* mutant. These FliG residues correspond to K300, R301, A302, D308, and D309 of *Vibrio*. We also expressed PomA L93C/PotB with I285C and L286C variants of *E. coli* FliG. I285 and L286 are hydrophobic residues located between R281 and D288 (Fig. S1B). Cells expressing only R281C FliG or L93C PomA lost motility in soft agar (Fig. S5), suggesting that these residues are critical for motor function. Cells expressing K89C PomA with wild-type FliG and Q280C, A282C, D288C or D289C FliG with wild-type PomA retained motility (Fig. S5). Cells co-expressing PomA K89C and FliG Q280C, A282C or D288C had reduced motility (Fig. S5).

We next tried to detect disulfide-crosslinked products between PomA and FliG. After oxidation with copper phenanthroline, we detected disulfide-crosslinked products of PomA K89C with FliG Q280C, R281C, A282C and D288C with anti-PomA or anti-FliG antibody (Fig. 4). The crosslinked products disappeared upon treatment with the reducing agent ?-mercaptoethanol (Fig. S6). In contrast, PomA L93C did not form disulfide crosslinks with either FliG I285C or L286C (Fig. 4).

**Fig. 4.**
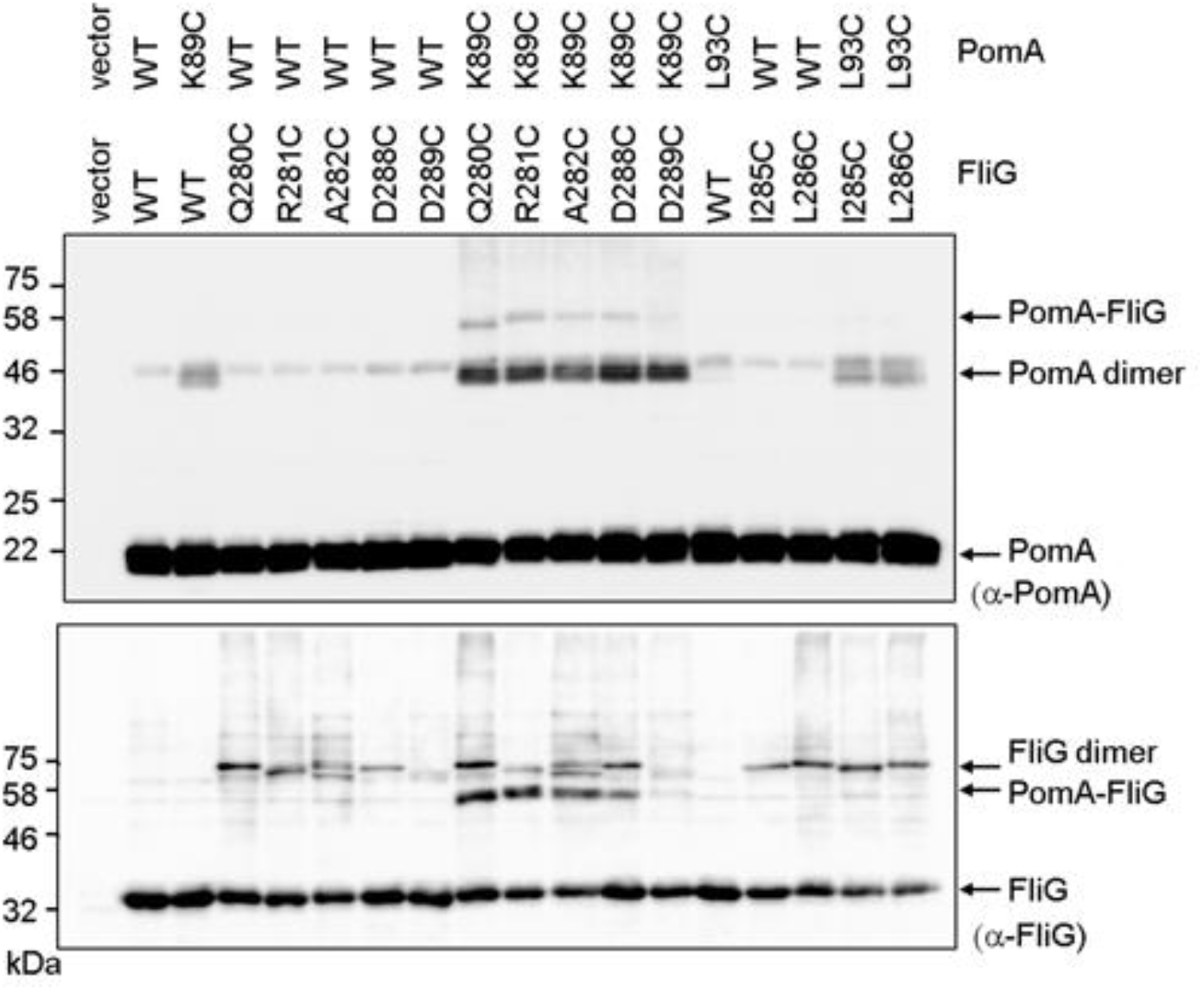
Disulfide-crosslinking between plasmid-borne *Vibrio* PomA and *E. coli* FliG. *Vibrio* PomA, chimeric PotB and *E. coli* FliG were co-expressed from plasmid pTSK170 in the *E. coli* Δ*motA*Δ*fliG* strain DFB245. The upper and lower panels show immunoblot images made using anti-PomA and anti-FliG antibodies, respectively.

### Effect of the conserved aspartate residue in the B subunit on crosslinking efficiency

Flagellar rotation is powered by conformational changes of the stator units that are driven by ion conduction. Therefore, we thought that the interaction pattern between PomA and FliG might be different in the presence and absence of Na+. We expressed PomA with various pBPA substitutions and PotB in *E. coli* Δ*motAB* cells and irradiated these cells with UV in the presence of Na+ or K+, followed by immunoblotting. There were no reproducible differences in crosslinking (Fig. S7). Next, we examined photo-crosslinking when the *p*BPA-substituted PomA proteins were co-expressed with PotB D24N, which has no ability to bind Na+ or support ion flow (30). The equivalent D32N substitution in *E. coli* (D33N in *Salmonella*) confers a dominant-negative effect on motility (28, 31, 50, 51). It has been speculated that aspartate to asparagine substitution mimics the protonated or Na+-bound state of the aspartyl residue. The crosslinking patterns with D24N PotB were similar to those seen with wild-type PotB (Fig. 5, S8), but the signal intensities of the crosslinked products with PotB D24N were stronger than those seen with wild-type PotB (Fig. 5).

**Fig. 5.**
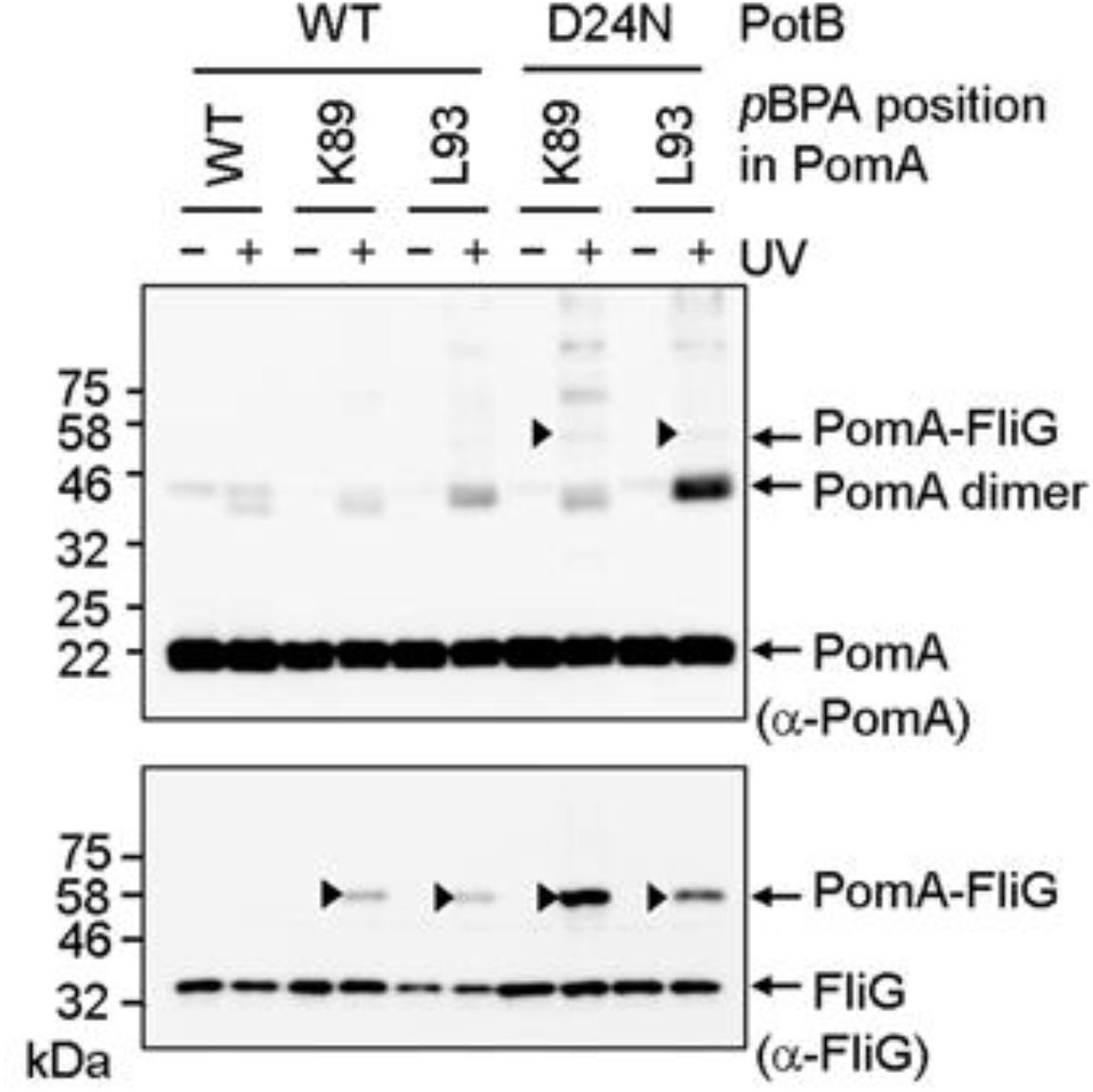
Photo-crosslinking between plasmid-borne *Vibrio* PomA and endogenous *E. coli* FliG in the presence of PotB D24N. *Vibrio* PomA and chimeric PotB were expressed from plasmid pYS3, and the amber suppressor tRNA and the mutated tyrosyl-tRNA synthase were expressed from plasmid pEVOL-pBpF in the *E. coli* Δ*motAB* strain RP6894. The upper and lower panels show immunoblot images of whole-cell lysates made using anti-PomA and anti-FliG antibodies, respectively. The crosslinked products are indicated by black arrowheads.

## Discussion

Previous genetic studies in *E. coli* and *Salmonella* showed that electrostatic interactions between the conserved charged residues of the stator A subunit and FliG in the rotor are important both for assembly of stator units into the motor and for torque generation (24, 34-37). Similar studies in *Vibrio alginolyticus* suggested that interactions in addition to the electrostatic interactions contribute to motor rotation (25, 38-40, 52). In this study, we used two different chemical crosslinking approaches to identify the residues in the *Vibrio* PomA subunit that are in close proximity to FliG, and the residues in *E. coli* FliG that are in close proximity to PomA. Crosslinked products were found in the isolated hook-basal body, indicating that crosslinking occurred in the intact motor. We also observed motility defects caused by photo-crosslinking, indicating that the crosslinking occurred, at least in part, within the motor.

Crosslinking was also observed between FliG and PomA in non-flagellated cells overexpressing soluble FliG, showing that stators freely diffusing in the cytoplasmic membrane interact with cytoplasmic FliG that is not assembled into a C-ring. Once the C-ring forms, FliG molecules in the C-ring do not exchange with cytoplasmic FliG monomers (53). Therefore, it is unlikely that the stator unit binds to the cytoplasmic FliG before the stator-FliG complex assembles into the motor. This result implies that a region around K89 is the first site in PomA to have access to FliG assembled into the rotor. Since PomA with *p*BPA replacing R88 in the chimeric PomA/PotB stator conferred a severe motility defect, we speculate that the stator complex containing PomA with *p*BPA replacing R88 assembles poorly into the motor. Previous studies too have suggested that the region around PomA R88 and K89 seems to be important for stator assembly into the motor rather than for torque generation (24, 25).

We found that the conserved motif RxxGΦΦxLE, which spans the region from PomA R88 to PomA E96, is important for motility (Fig. S1A). The motif contains a completely conserved G91 residue followed by two hydrophobic residues (Φ). Leucine (or less often isoleucine) at residue 95 is also highly conserved, and when it was replaced with *p*BPA, motility was abolished, suggesting that an aliphatic residue at this position is important for motor rotation. The hydrophobic residues F92 and L93 as well as the charged residues seem to contribute to the stator-rotor interaction. PomA with *p*BPA replacing L93 still supported motility, whereas the PomA L93C mutant did not support motility, suggesting that hydrophobicity at residue 93 is important for motor function. We showed the strong disulfide-crosslinking between *Vibrio* PomA K89C and *E. coli* FliG R281C or D288C. This result is consistent with the idea that electrostatic repulsion and attraction, respectively, between K89 in PomA and R281 and D288 contribute to torque generation. However, the residue at the position corresponding to K89 in PomA is Q in *E. coli* and *Salmonella* MotA, suggesting that these electrostatic interactions are not essential for motility in all cases. Overall, it seems that both electrostatic and hydrophobic interactions between the stator and rotor contribute to torque generation.

The three-dimensional structure at atomic resolution of the A subunit has been revealed by two independent groups (43, 44). The PDB structural data for MotA/MotB in *C. jejuni* were kindly supplied by Dr. Taylor before being available to the general public (Fig. 6). The residues corresponding to L76, I77, I80, A84, L95, N102 and F104, at which substitution with *p*BPA led to a complete loss of motility, were at positions internal to the MotA structure, suggesting that they are important for proper folding and stability. The residues corresponding to D85, R88, K89, G90, F92, L93 and E96 in PomA, which showed photo-crosslinking with FliG when replaced with *p*BPA, were arrayed on helices H1 and H2 and the H1-H2 linker. These residues are located on the most external and membrane-distal portion of MotA/PomA bound to the B subunit (Fig. S1C). The crosslinking data indicate that this surface, which contains the RxxGΦΦxLE motif, interacts closely with FliG and is important for stator assembly and motor rotation.

**Fig. 6.**
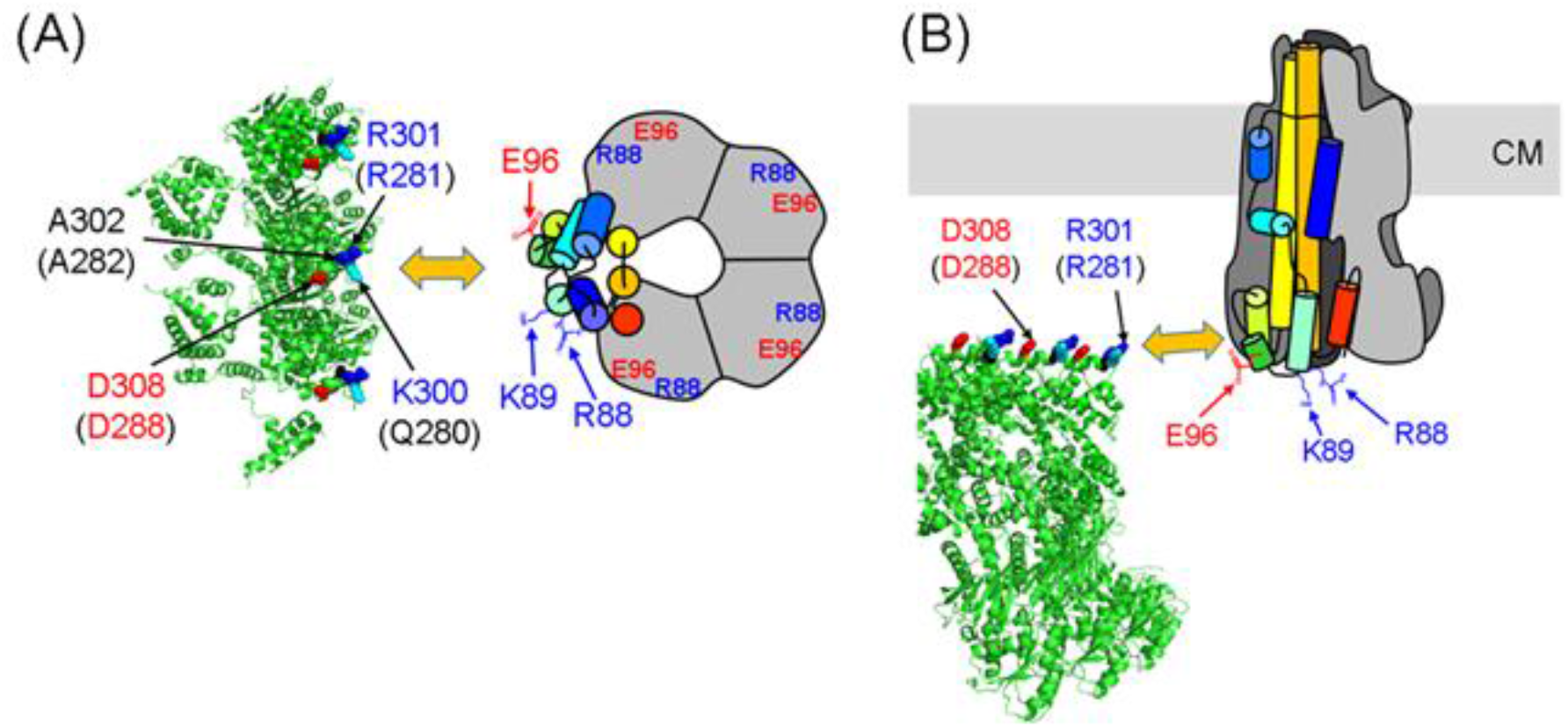
A model for the interaction between the C-ring and PomA from a top view (A) and a side view (B). The model incorporates a schematic of the PomA pentamer based on the cryo-EM structure (43). One PomA subunit is shown in rainbow color, and the other four PomA subunits are outlined in gray. The C-ring of *Vibrio alginolyticus* is based on a model previously reported (56). The positions of R88, K89 and E96 are indicated by arrows. FliG residues K300, R301 and A302 in *Vibrio,* corresponding to residues Q280, R281 and A282 in *E. coli* (the residue numbers for *E. coli* FliG are given in brackets), are shown as cyan, blue and red spheres, respectively. CM: cytoplasmic membrane.

The coupling ion for motility binds to an absolutely conserved aspartate residue in the B subunit. Substitution of this residue with asparagine, the D24N replacement in *Vibrio,* PomA may mimic the electrically neutral protonated state of MotA/MotB or the Na+-bound state of PomA/PomB. Therefore, the stator with the D24N mutation may not undergo conformational changes accompanied by Na+-binding or Na+-release from the aspartate residue. The chimeric PomA/PotB stator with the D24N variant of PotB formed the more crosslinked product with FliG than the stator containing the wild-type form of PotB. This result may imply that the D24N stator interacts with FliG more stable than the wild-type stator due to less movement in the region of the A subunit that interacts with FliG. However, this increased crosslinking may also reflect interactions of PomA/PotB D24N freely diffusing in the cytoplasmic membrane with cytoplasmic FliG.

Because we detected the interaction between *E. coli* FliG and *Vibrio* PomA within *E. coli* cells using the chimeric stator system, we cannot rule out the unlikely possibility that the residues in PomA and FliG that we have identified are not in close proximity in *Vibrio* cells. Eliminating this uncertainty would require comprehensive disulfide-crosslinking experiments with cysteine-substituted PomA and FliG in *Vibrio*.

Recently, a rotational gear model for stator function has been proposed by several groups (43, 44, 54). A similar rotational model has been reported for ExbB/ExbD in the Ton molecular motor. ExbB/ExbD shares a weak homology with MotA/MotB and PomA/PomB (43-47, 55). In the rotational gear model, the A subunit pentamer rotates around the B subunit dimer, and this rotation transmits torque to the rotor via interaction with the FliG ring. Based on our results, we can suggest which residues at the PomA (MotA) interface with FliG are involved in the interaction. We predicted that PomA K89 interacts with both R281 and D288 in *E. coli* FliG. We speculate that PomA residues R88 and K89 first interact with FliG D288 (D308 in *Vibrio* FliG) during assembly of a stator unit into the motor. This interaction activates ion conduction through the stator (24) and brings PomA E96 close to FliG R281 (R301 in *Vibrio*). Next, the A subunit begins to rotate in response to ion influx. Finally, PomA residues R88 and K89 repel the positive charge of FliG R281, and PomA E96 repels the negative charge of FliG D288 (D308 in *Vibrio*). As the stator rotation proceeds, the next PomA subunit interacts with the next FliG subunit on the C-ring (Fig. 6, S9). PomA residue L93 may also contribute to torque generation through hydrophobic interactions with FliG.

In summary, we provide the first biochemical evidence for close proximity of specific residues in the PomA/MotA stator with specific residues of FliG in the C-ring rotor. The presence of both charged and hydrophobic residues at these positions suggests that both electrostatic and hydrophobic interactions contribute to stator assembly and generation of rotation. These results provide insight into the fundamental molecular mechanisms of stator assembly around the rotor and torque generation within the flagellar motor.

## Materials and Methods

### Bacterial strains and plasmids

The bacterial strains and plasmids are listed in Table S1. *E. coli* was cultured in LB broth [1% (w/v) bactotryptone, 0.5% (w/v) yeast extract, 0.5% (w/v) NaCl], and TG broth [1% (w/v) bactotryptone, 0.5% (w/v) NaCl, 0.5% (w/v) glycerol]. Chloramphenicol was added to a final concentration of 25 µg/mL for *E. coli*. Ampicillin was added to a final concentration of 100 µg/mL for *E. coli*.

### Swimming assay in semi-soft agar

*E. coli* RP6894 cells harboring both pYS3 and pEVOL-pBpF, or *E. coli* DFB245 cells harboring pTSK170, were plated on LB agar with antibiotics. A single colony from the LB plate was inoculated onto TG agar plates [TG containing 0.3% (w/v) bactoagar and 0.02% (w/v) arabinose] and incubated at 30 °C for 24 hrs. For the dominant-negative experiment with PotB D24N, *E. coli* RP437 cells expressing PomA/PotB D24N were pre-cultured in LB broth with 0.2% (w/v) arabinose overnight at 30 °C, then 1 μL overnight cell culture was inoculated in TG semi-soft agar [TG containing 0.3% (w/v) bactoagar and 0.2% (w/v) arabinose] at 30 °C for 7 hrs.

### Photo-crosslinking experiment

*E. coli* cells harboring two different plasmids, pEVOL-pBpF and a pBAD24-based plasmid, were cultured in TG broth containing 1 mM *p*-benzoyl-L-phenylalanine (*p*BPA) (Bachem AG, Switzerland) at 30 °C for 2 hrs from an initial OD600 of 0.1. Arabinose was added to a final concentration of 0.02% (w/v) to express mutated tyrosyl-tRNA synthase, amber suppressor tRNA, and PomA/PotB and/or FliG, and further cultivated for 4 hrs. The cells were collected by centrifugation (3,400 × *g*, 5 min), resuspended in PBS buffer [137 mM NaCl, 2.7 mM KCl, 10 mM Na2HPO4, 1.76 mM KH2PO4], re-collected by centrifugation, and resuspended in PBS buffer. In the experiment shown in Fig. S7, we used sodium buffer [20 mM Tris-HCl pH 8.0, 150 mM NaCl] or potassium buffer [20 mM Tris-HCl pH 8.0, 150 mM KCl] instead of PBS. UV irradiation was performed with a B-100AP UV lamp (Analytik Jena US, Upland, CA, USA) for 5 min. The cells were collected by centrifugation (3,400 × *g*, 5 min) and resuspended in sodium dodecyl sulfate (SDS) loading buffer [62.5 mM Tris-HCl pH 6.8, 2% (w/v) SDS, 10% (w/v) glycerol, 0.01% (w/v) bromophenol blue] containing 5% (v/v) β-mercaptoethanol. The samples were separated by SDS-polyacrylamide gel electrophoresis (SDS-PAGE) and transferred to poly-vinylidene di-fluoride membrane. The proteins were detected using rabbit anti-PomA antibody and rabbit anti-*Salmonella* FliG antibody (a gift from Dr. Minamino at Osaka University). For the experiment shown in Figure 3, rabbit anti-*E. coli* FliG antibody (a gift from Dr. Blair at University of Utah) was used. The rabbit anti-*Salmonella* FliG antibody cross-reacts with *E. coli* FliG and can detect *E. coli* FliG at chromosomal levels.

### Purification of the hook-basal body complex

The hook-basal body complex was isolated as described previously, with several modifications (8). After UV irradiation for photo-crosslinking, the cells were suspended in 100 μL sucrose solution [0.5 M sucrose, 50 mM Tris-HCl pH 8.0]. EDTA, lysozyme and DNase were added to final concentrations of 10 mM, 1 mg/mL and 1 mg/mL, respectively. The suspension was left on ice for 30 min, and then spheroplasts were lysed by adding Triton X-100 and MgSO4 to final concentrations of 1% (w/v) and 15 mM, respectively. The lysate was then incubated on ice for 1 hr. After removal of the cell debris by centrifugation at 15,000 × *g* for 10 min, hook-basal bodies were precipitated by centrifugation at 60,000 × *g* for 60 min. The precipitate was resuspended in the SDS-loading buffer.

### Disulfide-crosslinking experiment

*E. coli* DFB245 strain harboring the pTSK170 plasmid was cultured in TG broth containing arabinose at a final concentration of 0.02% (w/v) at 30 °C for 5 hrs from an initial OD600 of 0.05. The cells were collected by centrifugation (3,400 × *g*, 5 min), resuspended in PBS buffer, collected by centrifugation, and resuspended in PBS buffer. To form the disulfide crosslink, 1 mM copper phenanthroline was added to the cell suspension, which was then incubated for 5 min. To stop the crosslinking, 3 mM N-methylmaleimide was added to the cell suspension and further incubated for 5 min. The cells were then collected by centrifugation (3,400 × *g*, 5 min) and suspended in the SDS-loading buffer without β-mercaptoethanol. The procedure for SDS-PAGE and immunoblotting was the same as that used in the photo-crosslinking experiment.

## Acknowledgments

We thank Dr. Peter G. Schultz for the kind gift of pEVOL-pBpF for photo-crosslinking, Dr. Tohru Minamino for the kind gift of rabbit anti-*Salmonella* FliG antibody, Dr. David F. Blair for the kind gift of rabbit anti-*E. coli* FliG antibody, Dr. T. Yorimitsu for the construction of plasmid pTY801 and Dr. Nicholas M.I. Taylor for the kind gift of PDB data for MotA/MotB (6YKM, 6YKP and 6YKR) before it was available to the public. We thank Dr. Mike Manson for critical reading for the manuscript, and K. Maki, Y. Kawase and A. Abe for technical support. This work was supported in part by JSPS KAKENHI Grant Numbers 18K07108 (to H.T.), 16H04774 and 18K19293 (to S.K.), and 20H03220 (to M.H.), and MEXT KAKENHI Grant Number 20H04864 (to H.T.).

---

**Fig. S1.**
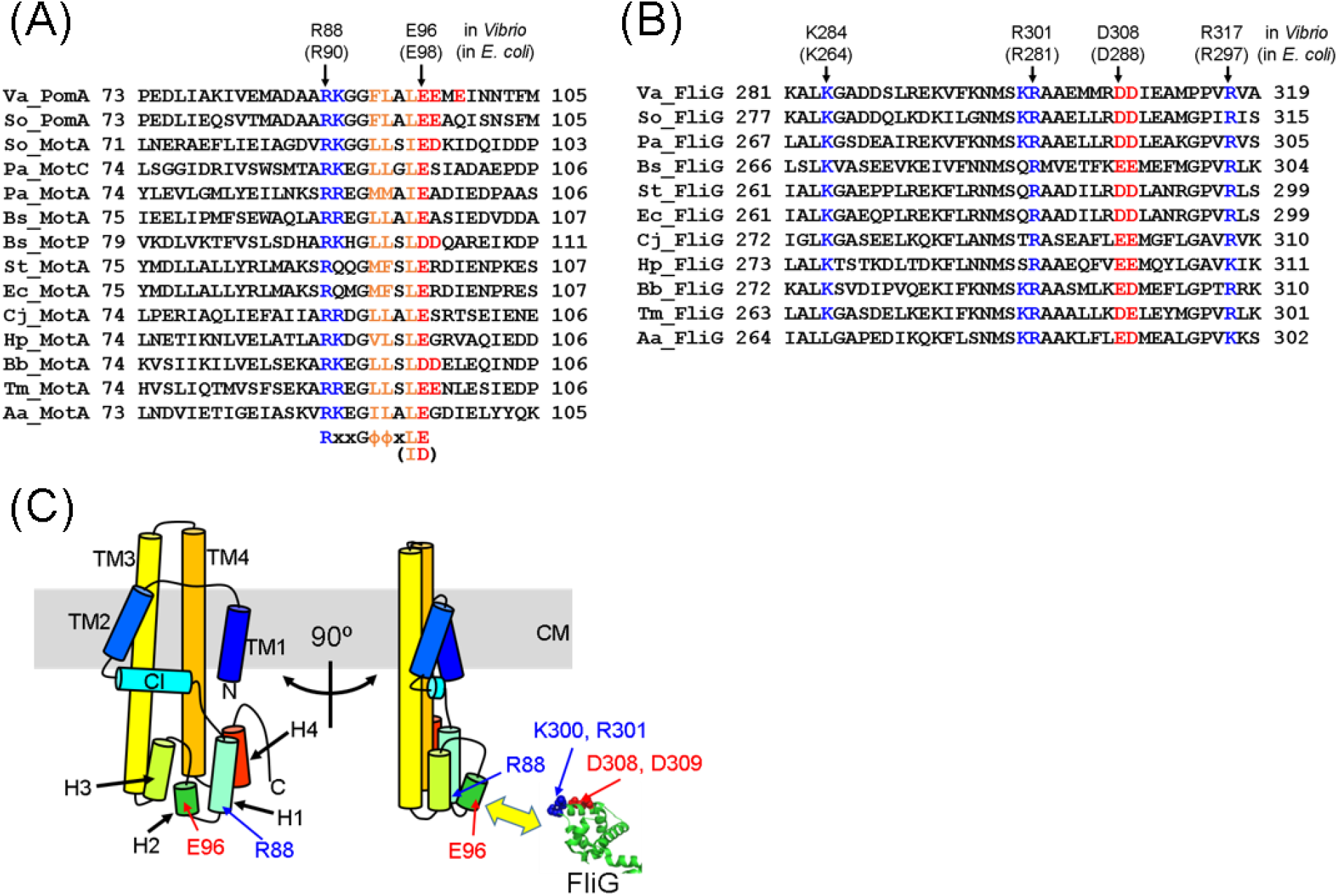
(A, B) Alignments of the amino acid sequences of PomA and MotA (A) and FliG(B) from various species. The charged residues thought to be important for the motor function were shown in blue (positive) and red (negative). The conserved hydrophobic residues in the RxxGΦΦxLE motif were shown in orange. Abbreviations: Φ, hydrophobic residues; Va, *Vibrio alginolyticus* VIO5; So, *Shewanella oneidensis* MR-1; Pa, *Pseudomonas aeruginosa* PAO1; Bs, *Bacillus subtilis* subsp. *subtilis* 168; St, *Salmonella enterica* subsp. *enterica* serovar *Typhimurium* LT2; Ec, *Escherichia coli* K-12 MG1655; Cj, *Campylobacter jejuni* subsp. *jejuni* NCTC 11168; Hp, *Helicobacter pylori* 26695; Bb, *Borreliella burgdorferi* B31; Tm, *Thermotoga maritima* MSB8; Aa, *Aquifex aeolicus* VF5. (C) The schematic drawing of the PomA monomer based on the cryo-EM structure of *C. jejuni* MotA/MotB complex (43). It was shown as a cartoon model in rainbow color. The mutations were introduced in H1, H1-H2 linker, H2 and H2-H3 linker. The C-terminal region of FliG from *A.aeolicus* was shown as a cartoon model (PDB ID: 3HJL). The important charged residues corresponding to *Vibrio* FliG K300, R301, D308 and D309, were shown in blue (positive) and red (negative) spheres. TM1∼TM4: transmembrane segments, CI: cytosolic interface helix, H1∼H4: cytosolic helices.

**Fig. S2.**
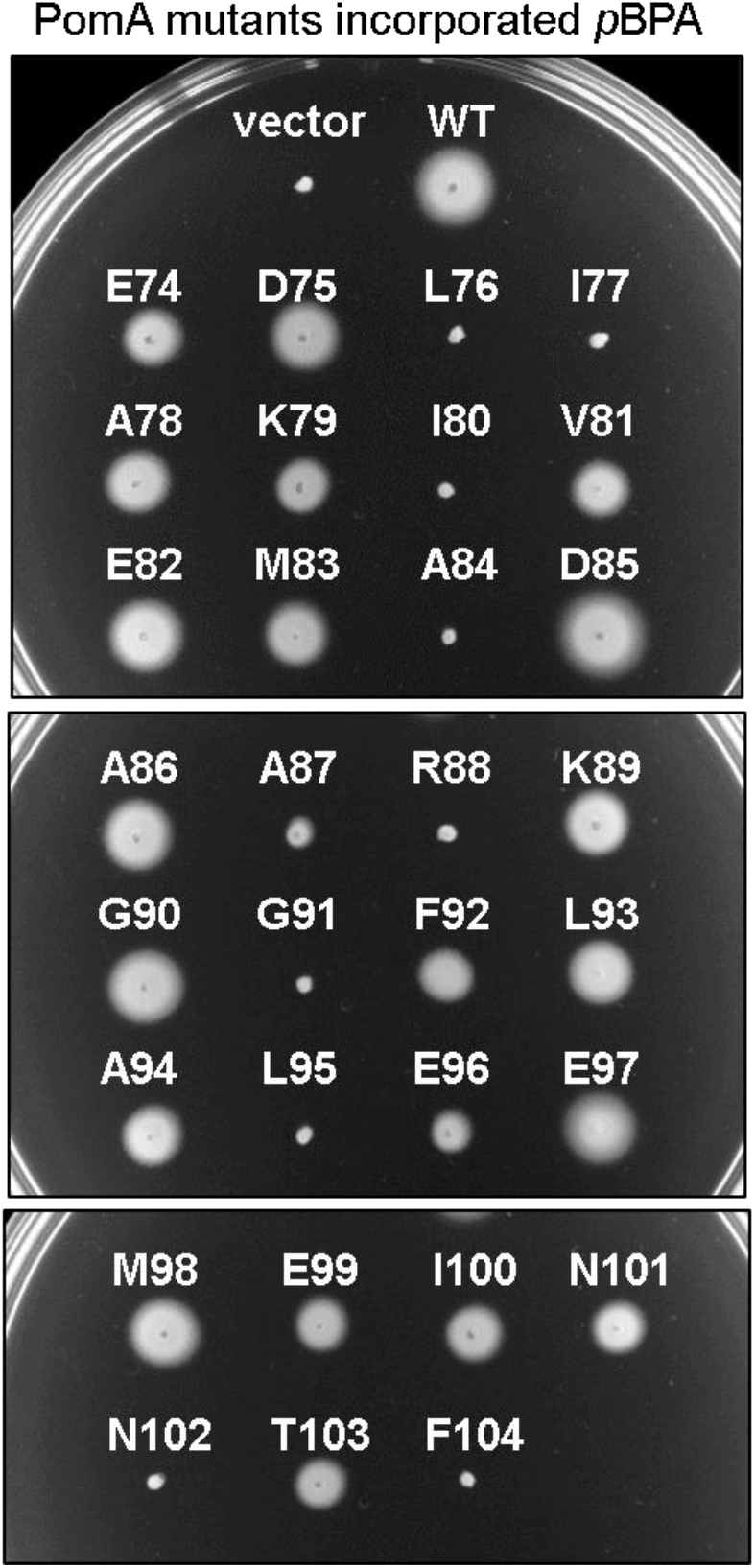
Motility of *E. coli* Δ*motAB* cells expressing *p*BPA-incorporated *Vibrio* PomA and chimeric PotB in a soft-agar plate. The cells were inoculated in TG 0.3% (w/v) bactoagar with 0.02% (w/v) arabinose plate at 30 °C for 24 hrs. The *E. coli* Δ*motAB* strain is RP6894. The vector plasmid is pBAD24. PomA/PotB were expressed from pYS3 that harbors *pomA* and *potB* genes in pBAD24 backbone. *p*BPA-incorporation into an amber codon was carried out by the amber suppressor tRNA and the mutated tyrosyl-tRNA synthase expressed from pEVOL-pBpF.

**Fig. S3.**
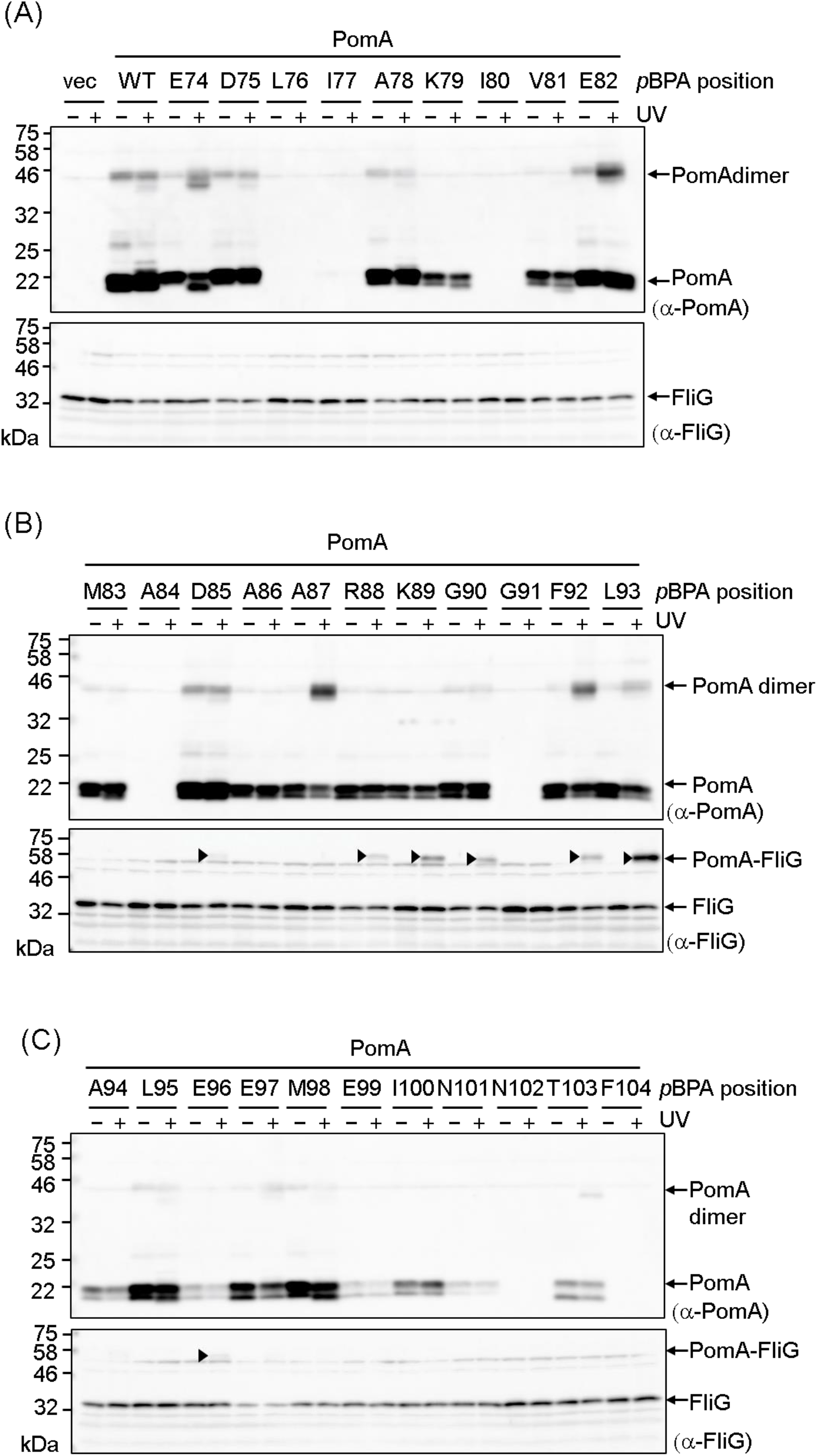
Protein expression of *p*BPA-introduced, plasmid-borne *Vibrio* PomA and photo-crosslinking between those PomA and endogenous *E. coli* FliG (A-C). SDS-PAGE samples were prepared from whole cell lysates. *Vibrio* PomA and chimeric PotB were expressed from plasmid pYS3, and the amber suppressor tRNA and the mutated tyrosyl-tRNA synthase were expressed from plasmid pEVOL-pBpF, in the *E. coli* Δ*motAB* strain, RP6894. Upper and lower panels showed immunoblot images by using anti-PomA and anti-FliG antibodies, respectively. The crosslinked products were marked by black arrow head.

**Fig. S4.**
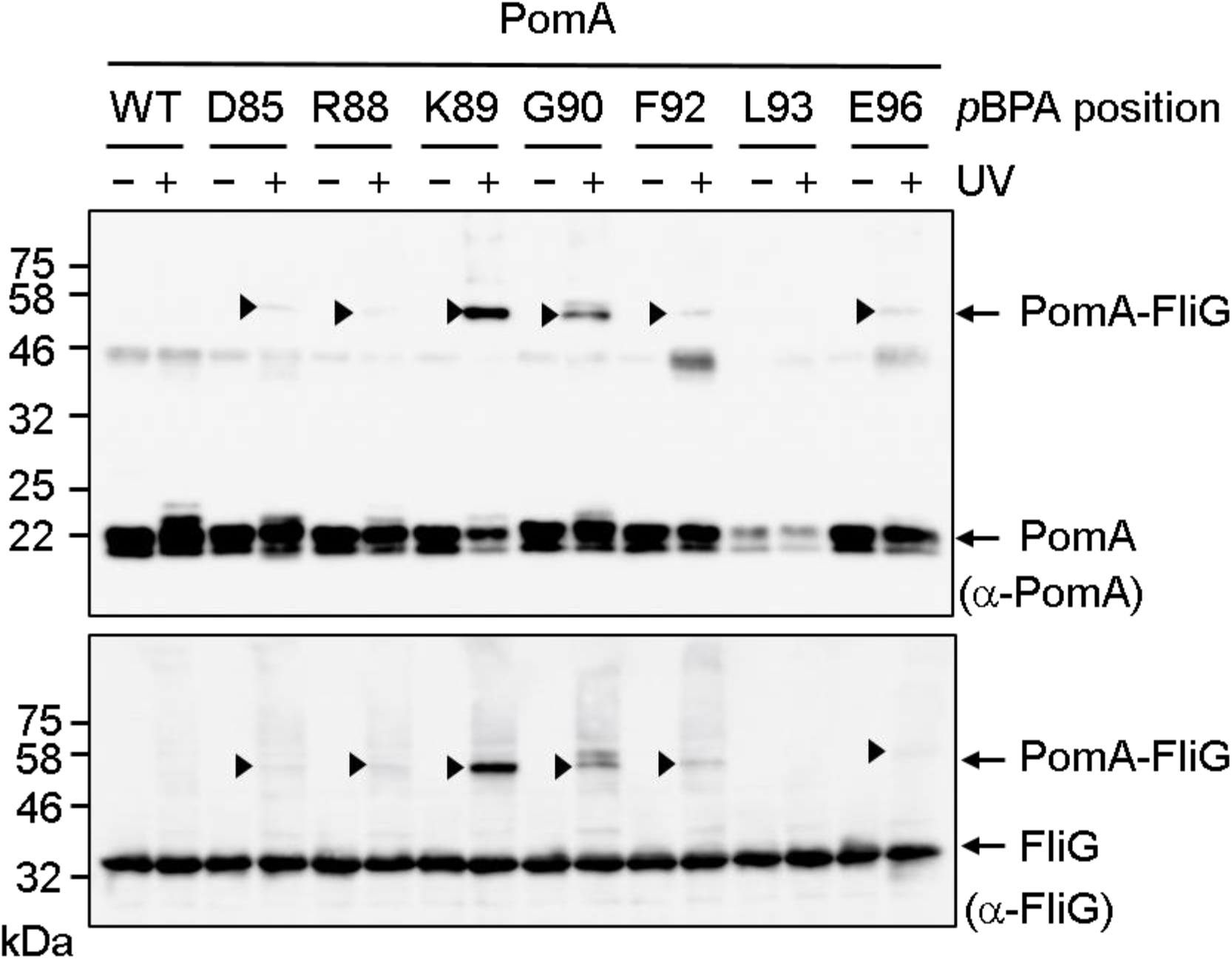
Photo-crosslinking between plasmid-borne *Vibrio* PomA and *E. coli* FliG in the *E. coli* Δ*flhDC* strain, RP3098. *E. coli* FliG, *Vibrio* PomA, chimeric PotB and *E. coli* FliG were co-expressed from plasmid pTSK170, and the amber suppressor tRNA and the mutated tyrosyl-tRNA synthase were expressed from plasmid pEVOL-pBpF. Upper and lower panels showed immunoblot images by using anti-PomA and anti-FliG antibodies, respectively. The crosslinked products were marked by black arrow head.

**Fig. S5.**
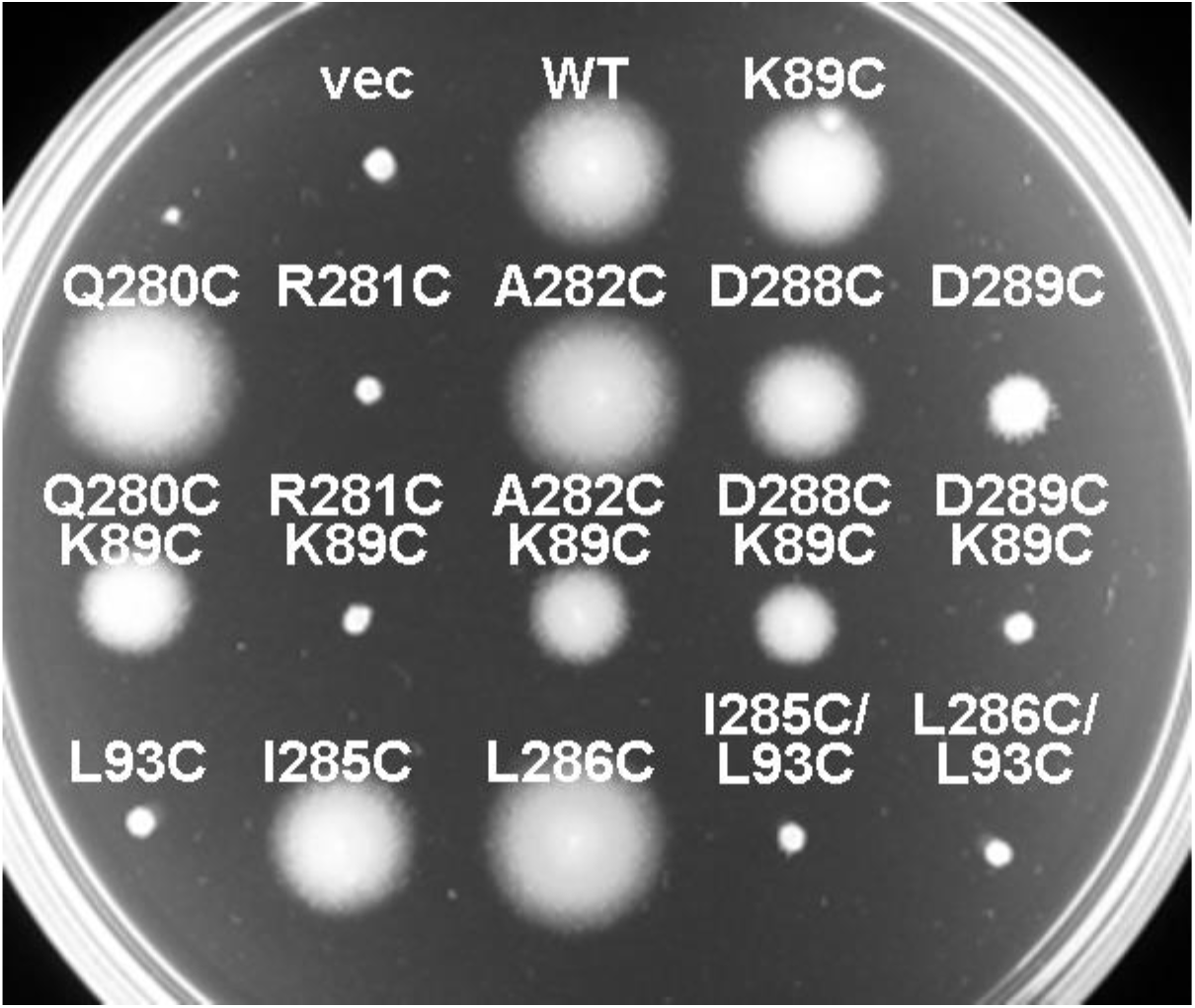
Motility of *E. coli* Δ*motA*Δ*fliG* cells expressed *Vibrio* PomA, and chimeric PotB and *E. coli* FliG in a soft-agar plate. The cells were inoculated in TG 0.3% (w/v) bactoagar with 0.02% (w/v) arabinose plate, and incubated at 30 °C for 24 hrs. The *E. coli* Δ*motA*Δ*fliG* strain is DFB245. The vector plasmid is pBAD24. PomA, PotB and FliG were expressed from pTSK170, in which *pomA, potB* and *fliG* genes were cloned into pBAD24.

**Fig. S6.**
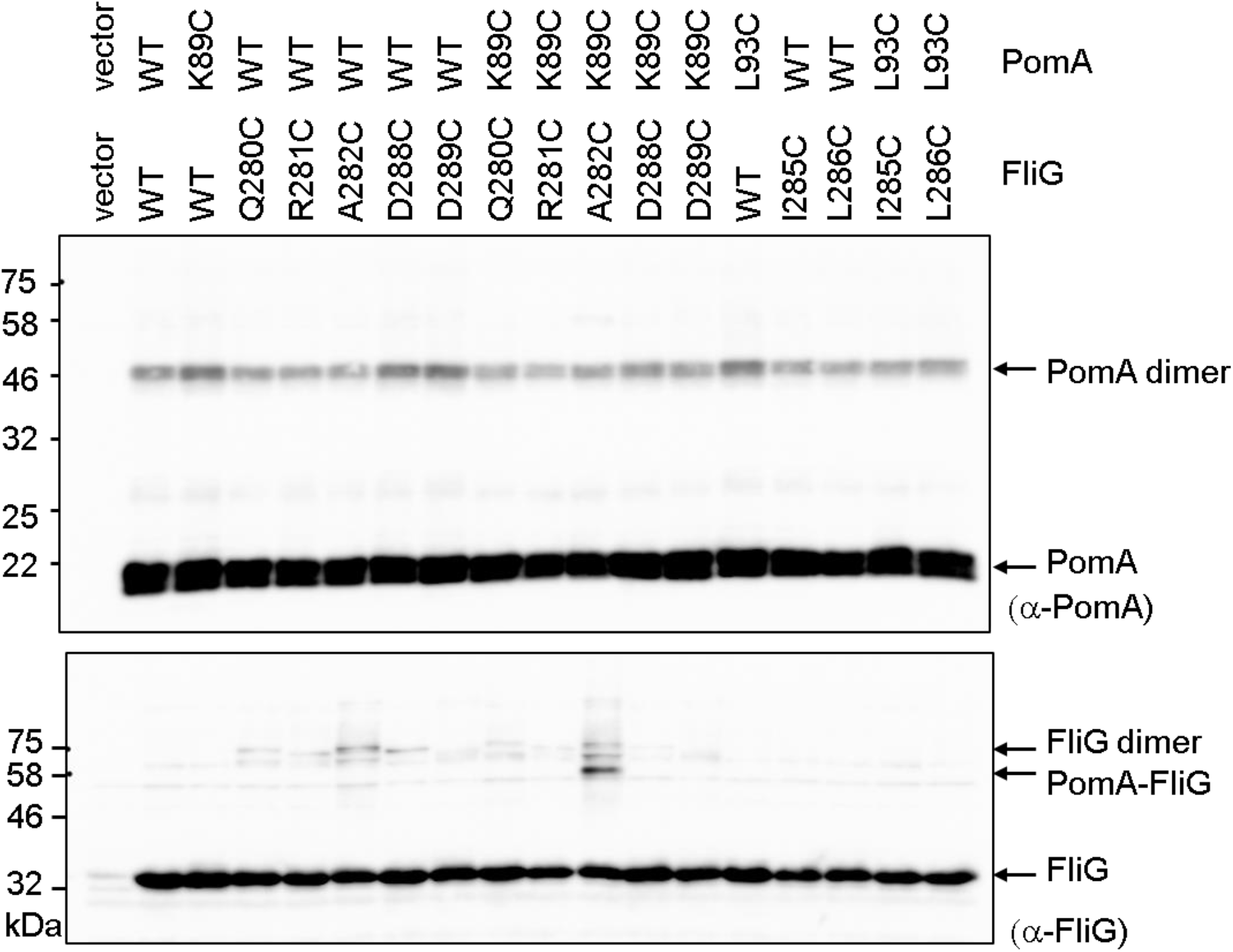
Immunoblotting of the disulfide crosslinked samples with reduced treatment by β-mercaptoethanol. *Vibrio* PomA, chimeric PotB and *E. coli* FliG were expressed from plasmid pTSK170, in the *E. coli* Δ*motA*Δ*fliG* strain, DFB245. Upper and lower panels showed immunoblot images by using anti-PomA and anti-FliG antibodies, respectively. A small amount of the crosslink product of FliG A282C/PomA K89C was detected even upon treatment with reducing agent.

**Fig. S7.**
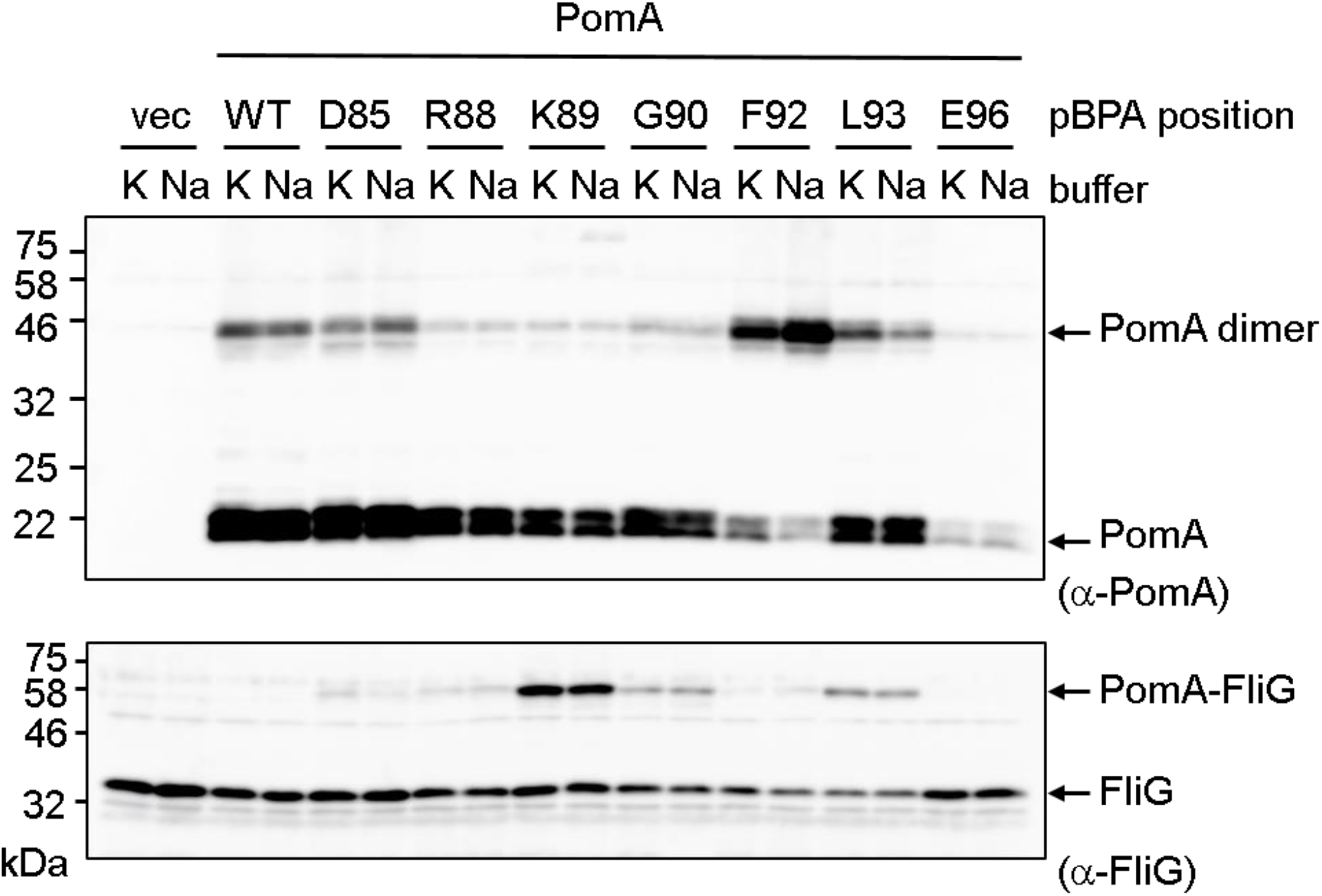
Photo-crosslinking between plasmid-borne *Vibrio* PomA and endogenous *E. coli* FliG in the presence of sodium buffer or potassium buffer. *Vibrio* PomA and chimeric PotB were expressed from plasmid pYS3, and the amber suppressor tRNA and the mutated tyrosyl-tRNA synthase were expressed from plasmid pEVOL-pBpF, into the *E. coli* Δ*motAB* strain, RP6894. Upper and lower panels showed immunoblot images by using anti-PomA and anti-FliG antibodies, respectively.

**Fig. S8.**
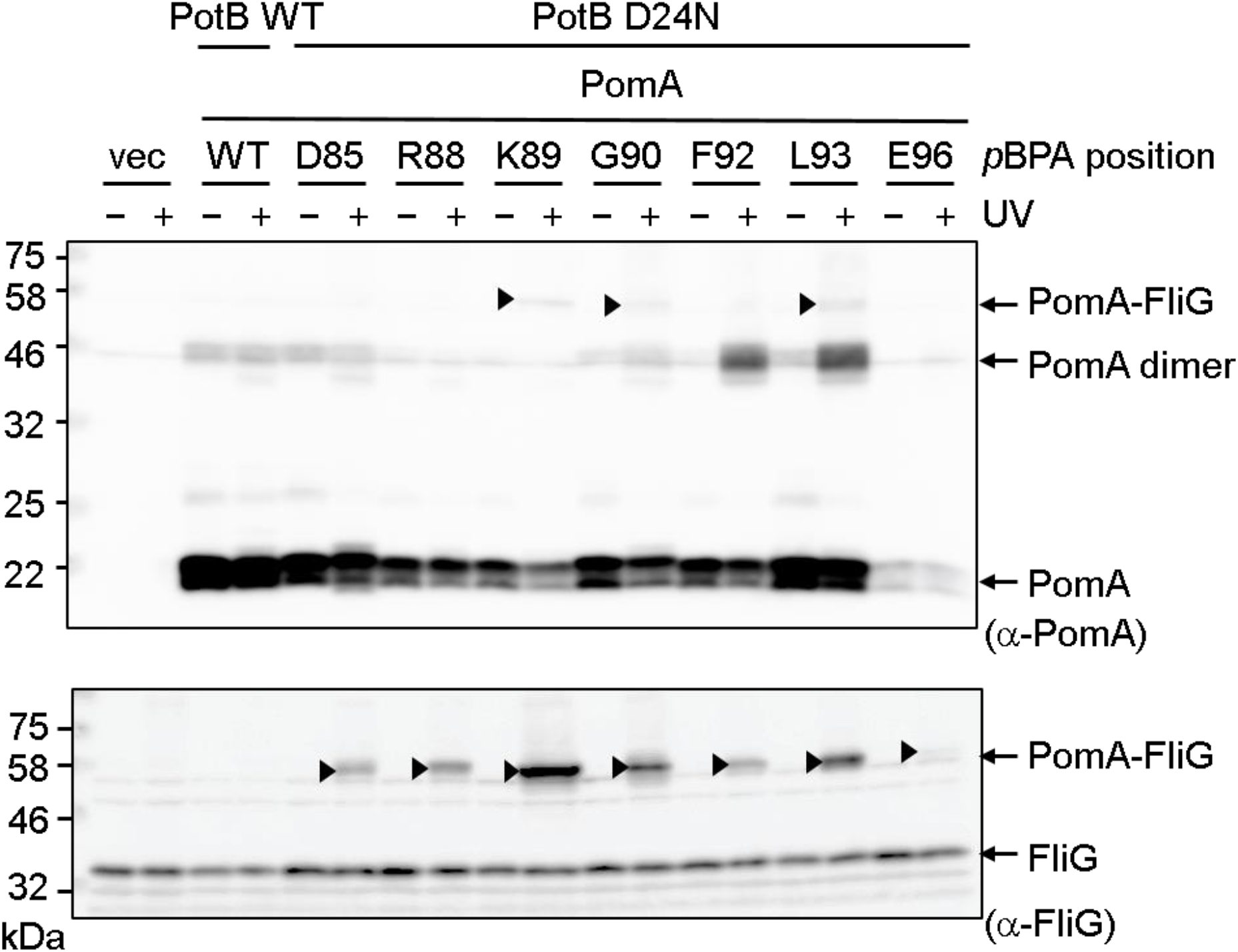
Photo-crosslinking between plasmid-borne *Vibrio* PomA and endogenous *E. coli* FliG in the background of PotB D24N. *Vibrio* PomA and chimeric PotB were expressed from plasmid pYS3, and the amber suppressor tRNA and the mutated tyrosyl-tRNA synthase were expressed from plasmid pEVOL-pBpF, into the *E. coli* Δ*motAB* strain, RP6894. Upper and lower panels showed immunoblot images by using anti-PomA and anti-FliG antibodies, respectively. The crosslinked products were marked by black arrow head.

**Fig. S9.**
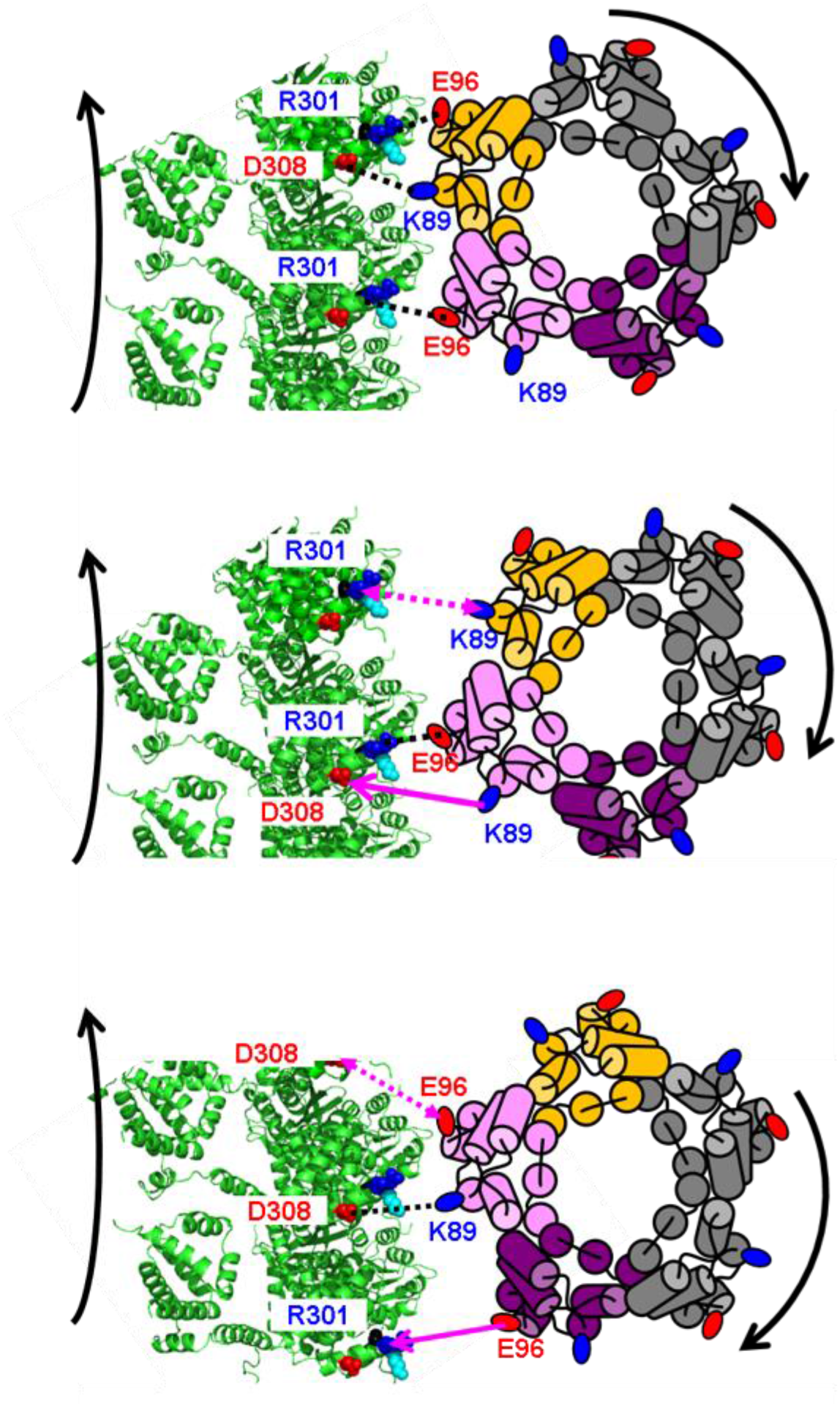
The interaction model between C-ring and PomA. The schematic diagram of the PomA pentamer based on the cryo-EM structure (43) was shown from the top view. The C-ring model of *Vibrio alginolyticus* was based on the model previously reported (56) is shown from the top view. The positions of PomA K89 and E96 were shown in blue and red circles, respectively. FliG R301 and D308 in *Vibrio* corresponding to R281 and D288 in *E. coli* were shown by space filling residues.

**Table S1.** Bacterial strains and plasmids used in this study

| Strain or plasmid | Genotype or description | Reference or source |
| --- | --- | --- |
| *E. coli* |  |  |
| DH5α | F<sup>-</sup> Φ80d*lacZ* ΔM15 $\Delta$(*lacZYA-argF*)U169 *deoR recA1 endA1 hsdR17*(r$\kappa^-$, m$\kappa^+$) *phoA supE44* $\lambda^-$ *thi-1 gyrA96 relA1* | (57) |
| RP437 | Wild-type strain | (58) |
| RP6894 | *motA* and *motB* null strain | (59) |
| RP3098 | *flhD* and *flhC* null strain | (60) |
| DFB245 | *motA* and *fliG* null strain | (34) |
| DFB225 | *fliG* null strain | (10) |
| Plasmids |  |  |
| pBAD24 | pBR322-derived vector, *araBAD* promoter, Amp<sup>r</sup> | (61) |
| pBAD33 | pACYC-derived vector, *araBAD* promoter, Cm<sup>r</sup> | (61) |
| pYS3 | *Vibrio pomA* and chimeric *potB* in pBAD24 | (40) |
| pTSK170 | *E. coli fliG*, *Vibrio pomA* and chimeric *potB* in pBAD24 | This study |
| pTY801 | *E. coli fliG* in pBAD24 | This study |
| pEVOL-pBpF | Plasmid for the incorporation of photo-reactive amino acid, *pBPA*, into the amber codon. | (48) |
| Amp<sup>r</sup>, ampicillin-resistant; Cm<sup>r</sup>, chloramphenicol-resistant. |  |  |

## References

1. Stewart AG, Laming EM, Sobti M, Stock D (2014) Rotary ATPases--dynamic molecular machines. Curr Opin Struct Biol 25:40–48.

2. Nakanishi A, Kishikawa JI, Mitsuoka K, Yokoyama K (2019) Cryo-EM studies of the rotary H+-ATPase/synthase from Thermus thermophilus. Biophy Physicobiol 16:140–146.

3. Macnab RM (2003) How bacteria assemble flagella. Ann Rev Microbiol 57:77–100.

4. Terashima H, Kojima S, Homma M (2008) Flagellar motility in bacteria structure and function of flagellar motor. Int Rev Cell Mol Biol 270:39–85.

5. Morimoto YV, Minamino T (2014) Structure and function of the bi-directional bacterial flagellar motor. Biomolecules 4:217–234.

6. Takekawa N, Imada K, Homma M (2020) Structure and energy-conversion mechanism of bacterial Na+-driven flagellar motor. Trends Microbiol 28:719–731.

7. Ueno T, Oosawa K, Aizawa SI (1992) M-Ring, S-ring and proximal rod of the flagellar basal body of Salmonella-typhimurium are composed of subunits of a single protein, FliF. J Mol Biol 227:672–677.

8. Francis NR, Sosinsky GE, Thomas D, Derosier DJ (1994) Isolation, characterization and structure of bacterial flagellar motors containing the switch complex. J Mol Biol 235:1261–1270.

9. Yamaguchi S, Aizawa S, Kihara M, Isomura M, Jones CJ, Macnab RM (1986) Genetic evidence for a switching and energy-transducing complex in the flagellar motor of Salmonella typhimurium. J Bacteriol 168:1172–1179.

10. Lloyd SA, Tang H, Wang X, Billings S, Blair DF (1996) Torque generation in the flagellar motor of Escherichia coli: Evidence of a direct role for FliG but not for FliM or FliN. J Bacteriol 178:223–231.

11. Thomas DR, Francis NR, Xu C, DeRosier DJ (2006) The three-dimensional structure of the flagellar rotor from a clockwise-locked mutant of Salmonella enterica serovar Typhimurium. J Bacteriol 188:7039–7048.

12. Dean GD, Macnab RM, Stader J, Matsumura P, Burks C (1984) Gene sequence and predicted amino acid sequence of the motA protein, a membrane-associated protein required for flagellar rotation in Escherichia coli. J Bacteriol 159:991–999.

13. Stader J, Matsumura P, Vacante D, Dean GE, Macnab RM (1986) Nucleotide sequence of the Escherichia coli MotB gene and site-limited inccorporation of its product into the cytoplasmic membrane. J Bacteriol 166:244–252.

14. Kojima S, Blair DF (2004) Solubilization and purification of the MotA/MotB complex of Escherichia coli. Biochemistry 43:26–34.

15. Asai Y, Kojima S, Kato H, Nishioka N, Kawagishi I, Homma M (1997) Putative channel components for the fast-rotating sodium-driven flagellar motor of a marine bacterium. J Bacteriol 179:5104–5110.

16. Sato K, Homma M (2000) Multimeric structure of PomA, the Na+-driven polar flagellar motor component of Vibrio alginolyticus. J Biol Chem 275:20223–20228.

17. De Mot R, Vanderleyden J (1994) The C-terminal sequence conservation between OmpA-related outer membrane proteins and MotB suggests a common function in both gram-positive and gram-negative bacteria, possibly in the interaction of these domains with peptidoglycan. Mol Microbiol 12:333–334.

18. Kojima S, Imada K, Sakuma M, Sudo Y, Kojima C, Minamino T, Homma M, Namba K (2009) Stator assembly and activation mechanism of the flagellar motor by the periplasmic region of MotB. Mol Microbiol 73:710–718.

19. Zhu S, Takao M, Li N, Sakuma M, Nishino Y, Homma M, Kojima S, Imada K (2014) Conformational change in the periplamic region of the flagellar stator coupled with the assembly around the rotor. Proc Natl Acad Sci USA 111:13523–13528.

20. Leake MC, Chandler JH, Wadhams GH, Bai F, Berry RM, Armitage JP (2006) Stoichiometry and turnover in single, functioning membrane protein complexes. Nature 443:355–358.

21. Reid SW, Leake MC, Chandler JH, Lo C-J, Armitage JP, Berry RM (2006) The maximum number of torque-generating units in the flagellar motor of Escherichia coli is at least 11. Proc Natl Acad Sci USA 103:8066–8071.

22. Kojima S, Takao M, Almira G, Kawahara I, Sakuma M, Homma M, Kojima C, Imada K (2018) The helix rearrangement in the periplasmic domain of the flagellar stator B subunit activates peptidoglycan binding and ion influx. Structure 26:590–598.

23. Zhu S, Nishikino T, Takekawa N, Terashima H, Kojima S, Imada K, Homma M, Liu J (2020) In situ structure of the Vibrio polar flagellum reveals a distinct outer membrane complex and its specific interaction with the stator. J Bacteriol 202:e00592–19.

24. Morimoto YV, Nakamura S, Hiraoka KD, Namba K, Minamino T (2013) Distinct roles of highly conserved charged residues at the MotA-FliG interface in bacterial flagellar motor rotation. J Bacteriol 195:474–481.

25. Takekawa N, Kojima S, Homma M (2014) Contribution of many charged residues at the stator-rotor interface of the Na+-driven flagellar motor to torque generation in Vibrio alginolyticus. J Bacteriol 196:1377–1385.

26. Blair DF, Berg HC (1990) The MotA protein of E. coli is a proton-conducting component of the flagellar motor. Cell 60:439–449.

27. Sato K, Homma M (2000) Functional reconstitution of the Na+-driven polar flagellar motor component of Vibrio alginolyticus. J Biol Chem 275:5718–5722.

28. Zhou J, Sharp LL, Tang HL, Lloyd SA, Billings S, Braun TF, Blair DF (1998) Function of protonatable residues in the flagellar motor of Escherichia coli: a critical role for Asp 32 of MotB. J Bacteriol 180:2729–2735.

29. Sudo Y, Kitade Y, Furutani Y, Kojima M, Kojima S, Homma M, Kandori H (2009) Interaction between Na+ ion and carboxylates of the PomA-PomB stator unit studied by ATR-FTIR spectroscopy. Biochemistry 48:11699–11705.

30. Onoue Y, Iwaki M, Shinobu A, Nishihara Y, Iwatsuki H, Terashima H, Kitao A, Kandori H, Homma M (2019) Essential ion binding residues for Na+ flow in stator complex of the Vibrio flagellar motor. Sci Rep 9:11216.

31. Kojima S, Blair DF (2001) Conformational change in the stator of the bacterial flagellar motor. Biochemistry 40:13041–13050.

32. Mino T, Nishikino T, Iwatsuki H, Kojima S, Homma M (2019) Effect of sodium ions on conformations of the cytoplasmic loop of the PomA stator protein of Vibrio alginolyticus. J Biochem 166:331–341.

33. Tang H, Braun TF, Blair DF (1996) Motility protein complexes in the bacterial flagellar motor. J Mol Biol 261:209–221.

34. Zhou JD, Lloyd SA, Blair DF (1998) Electrostatic interactions between rotor and stator in the bacterial flagellar motor. Proc Natl Acad Sci USA 95:6436–6441.

35. Lloyd SA, Blair DF (1997) Charged residues of the rotor protein FliG essential for torque generation in the flagellar motor of Escherichia coli. J Mol Biol 266:733–744.

36. Zhou JD, Blair DF (1997) Residues of the cytoplasmic domain of MotA essential for torque generation in the bacterial flagellar motor. J Mol Biol 273:428–439.

37. Morimoto YV, Nakamura S, Kami-ike N, Namba K, Minamino T (2010) Charged residues in the cytoplasmic loop of MotA are required for stator assembly into the bacterial flagellar motor. Mol Microbiol 78:1117–1129.

38. Yorimitsu T, Sowa Y, Ishijima A, Yakushi T, Homma M (2002) The systematic substitutions around the conserved charged residues of the cytoplasmic loop of Na+-driven flagellar motor component PomA. J Mol Biol 320:403–413.

39. Yorimitsu T, Mimaki A, Yakushi T, Homma M (2003) The conserved charged residues of the C-terminal region of FliG, a rotor component of Na+-driven flagellar motor. J Mol Biol 334:567–583.

40. Yakushi T, Yang J, Fukuoka H, Homma M, Blair DF (2006) Roles of charged residues of rotor and stator in flagellar rotation: comparative study using H+-driven and Na+-driven motors in Escherichia coli. J Bacteriol 188:1466–1472.

41. Yonekura K, Maki-Yonekura S, Homma M (2011) The structure of the flagellar motor protein complex PomAB: Implications for the torque-generating conformation. J Bacteriol 193:3863–3870.

42. Takekawa N, Terahara N, Kato T, Gohara M, Mayanagi K, Hijikata A, Onoue Y, Kojima S, Shirai T, Namba K, Homma M (2016) The tetrameric MotA complex as the core of the flagellar motor stator from hyperthermophilic bacterium. Sci Rep 6:31526.

43. Santiveri M, Roa-Eguiara A, Kühne C, Wadhwa N, Hu H, Berg HC, Erhardt M, Taylor NMI (2020) Structure and function of stator units of the bacterial flagellar motor. Cell 183:244–257.

44. Deme JC, Johnson S, Vickery O, Aron A, Monkhouse H, Griffiths T, Hennell James R, Berks BC, Coulton JW, Stansfeld PJ, Lea SM (2020) Structures of the stator complex that drives rotation of the bacterial flagellum. Nat Microbiol 5:1553–1564.

45. Celia H, Noinaj N, Zakharov SD, Bordignon E, Botos I, Santamaria M, Barnard TJ, Cramer WA, Lloubes R, Buchanan SK (2016) Structural insight into the role of the Ton complex in energy transduction. Nature 538:60–65.

46. Maki-Yonekura S, Matsuoka R, Yamashita Y, Shimizu H, Tanaka M, Iwabuki F, Yonekura K (2018) Hexameric and pentameric complexes of the ExbBD energizer in the Ton system. eLife 7:e35419.

47. Celia H, Botos I, Ni X, Fox T, De Val N, Lloubes R, Jiang J, Buchanan SK (2019) Cryo-EM structure of the bacterial Ton motor subcomplex ExbB-ExbD provides information on structure and stoichiometry. Commun Biol 2:358.

48. Chin JW, Martin AB, King DS, Wang L, Schultz PG (2002) Addition of a photocrosslinking amino acid to the genetic code of Escherichia coli. Proc Natl Acad Sci USA 99:11020–11024.

49. Asai Y, Yakushi T, Kawagishi I, Homma M (2003) Ion-coupling determinants of Na+-driven and H+-driven flagellar motors. J Mol Biol 327:453–463.

50. Nakamura S, Kami-ike N, Yokota JP, Minamino T, Namba K (2010) Evidence for symmetry in the elementary process of bidirectional torque generation by the bacterial flagellar motor. Proc Natl Acad Sci USA 107:17616–17620.

51. Nakamura S, Minamino T, Kami-Ike N, Kudo S, Namba K (2014) Effect of the MotB(D33N) mutation on stator assembly and rotation of the proton-driven bacterial flagellar motor. Biophysics 10:35–41.

52. Fukuoka H, Yakushi T, Homma M (2004) Concerted effects of amino acid substitutions in conserved charged residues and other residues in the cytoplasmic domain of PomA, a stator component of Na+-driven flagella. J Bacteriol 186:6749–6758.

53. Fukuoka H, Yakushi T, Kusumoto A, Homma M (2005) Assembly of motor proteins, PomA and PomB, in the Na+-driven stator of the flagellar motor. J Mol Biol 351:707–717.

54. Chang Y. Zhang K, Carroll BL, Zhao X, Charon NW, Norris SJ, Motaleb MdA, Li C, Liu J (2020) Molecular Mechanism for Rotational Switching of the Bacterial Flagellar Motor. Nat Struct Mol Biol 27:1041–1047.

55. Celia H, Noinaj N, Buchanan SK (2020) Structure and Stoichiometry of the Ton Molecular Motor. Int J Mol Sci 21:375.

56. Nishikino T, Hijikata A, Miyanoiri Y, Onoue Y, Kojima S, Shirai T, Homma M (2018) Rotational direction of flagellar motor from the conformation of FliG middle domain in marine Vibrio. Sci Rep 8:17793.

57. Grant SG, Jessee J, Bloom FR, Hanahan D (1990) Differential plasmid rescue from transgenic mouse DNAs into Escherichia coli methylation-restriction mutants. Proc Natl Acad Sci USA 87:4645–4649.

58. Parkinson JS (1978) Complementation analysis and deletion mapping of Escherichia coli mutants defective in chemotaxis. J Bacteriol 135:45–53.

59. Braun TF, Poulson S, Gully JB, Empey JC, Van Way S, Putnam A, Blair DF (1999) Function of proline residues of MotA in torque generation by the flagellar motor of Escherichia coli. J Bacteriol 181:3542–3551.

60. Slocum MK, Parkinson JS (1983) Genetics of methyl-accepting chemotaxis proteins in Escherichia coli: organization of the tar region. J Bacteriol 155:565–577.

61. Guzman LM, Belin D, Carson MJ, Beckwith J (1995) Tight regulation, modulation, and high-level expression by vectors containing the arabinose pBAD promoter. J Bacteriol 177:4121–4130.

